# Daily orexinergic modulation of the rat superficial layers of the superior colliculus – implications for intrinsic clock activities in the visual system

**DOI:** 10.1101/2021.02.20.432093

**Authors:** Lukasz Chrobok, Jagoda Stanislawa Jeczmien-Lazur, Monika Bubka, Kamil Pradel, Aleksandra Klekocinska, Jasmin Daniela Klich, Amalia Ridla Rahim, Jihwan Myung, Mariusz Kepczynski, Marian Henryk Lewandowski

## Abstract

The orexinergic system delivers excitation for multiple brain centres to facilitate behavioural arousal, with its malfunction resulting in narcolepsy, somnolence, and notably, visual hallucinations. Since the circadian clock underlies the daily arousal, a timed coordination is expected between the orexin system and its target subcortical visual system, including the superior colliculus (SC). Here, we use a combination of electrophysiological, immunohistochemical, and molecular approaches across 24 h, together with the neuronal tract tracing methods in rodents to elucidate the daily coordination between the orexin system and the superficial layers of the SC. We find the daily orexinergic innervation onto the SC, coinciding with the daily silencing of spontaneous firing in this visual brain area. We identify autonomous daily and circadian expression of clock genes in the SC, which may underlie these day-night changes. Additionally, we establish the lateral hypothalamic origin of orexin innervation to the SC and that the SC neurons robustly respond to orexin A via OX_2_ receptor in both excitatory and GABA_A_ receptor-dependent inhibitory manners. Together, our evidence supports that the clock coordination between the orexinergic input and its response in the SC provides arousal-related excitation needed to detect sparse visual information during the behaviourally active phase.

## 1. INTRODUCTION

The orexinergic system originating in the lateral hypothalamus (LH) consists of two evolutionary conserved peptides: orexin A (OXA) and orexin B (OXB), which bind to two metabotropic receptors, named OX_1_ and OX_2_ receptors [1,2]. Neurons that synthesise orexins provide extensive innervation to multiple brain centres, delivering arousal-related information in a daily (under light-dark conditions, LD) and circadian time-dependent manner (under constant darkness, DD). The activity of orexinergic system confers wakefulness and promotes feeding [3,4]. As the central circadian clock, the suprachiasmatic nucleus of the hypothalamus (SCN) reciprocally connects with the orexinergic system and directs the profound day-to-night changes, orexins have been hypothesised to act as hands of the clock [5–8]. However, new findings question the omnipotent role of the SCN in the circadian control of the brain physiology and function, with several neuronal centres rhythmically expressing clock genes independently from the central clock [9–11]. The extent to which these local circadian clocks interact and play a role in the brain physiology is now an active area of research. Although some of the clock interactions have been known in the regulatory areas of the brain such as the hypothalamus [12], epithalamus (habenula: [13,14]), brainstem [15], forebrain circumventricular organs [16], and even the choroid plexus [17], it remains largely unexplored as to how the clock interactions extend to higher brain areas such as the visual system.

Growing evidence supports the functional link between the orexinergic and visual systems [18–24]. However, whether orexins influence the neurophysiology of the extra-geniculate pathway, including the superior colliculus (SC), has not been demonstrated so far. In rodents, the SC is a layered midbrain structure that orients the animal towards the stimuli of the outside world during the wake phase. It consists of the retinorecipient superficial layers, the deep layers implicated in motor control, and the intermediate layers associated with both motor functions and integration of the multisensory input [25–29]. In humans, impaired activity of the superficial layers of the SC has been shown to diminish saccadic eye movement, to cause attention deficits, and to promote visual hallucinations [30–32].

Last two decades were marked by the emergence of new drugs targeting the orexinergic system, with double orexin receptor antagonists (DORAs) showing benefits in improving sleep onset and sleep maintenance [33]. Thus, the discovery and description of orexin binding sites in the brain stands as both basic scientific and clinical achievement. Despite a distinct safety profile of DORAs, the most frequently reported side effects include somnolence, headache, and fatigue; in particular, the side effects also include visual hallucinations and nightmares, as if to suggest an internal misalignment of wake and sleep [34]. The SC is a component of subcortical visual system that has been associated with visual hallucinations in schizophrenia and dementia patients [31,35].

Importantly, all of these syndromes are also observed in narcoleptic patients with the malfunction of OX_2_ receptors [36].

The aim of this study is to evaluate whether the orexinergic system modulates neuronal activity of the visual layers of the SC in the time-of-day dependent manner, and if so, to identify a source (extrinsic vs. intrinsic) of this temporal variation. Here, we find compelling evidence for the robust modulatory action of orexins upon the neuronal activity in the retinorecipient superficial layers of the SC. These neurophysiological effects were not exclusively excitatory but also inhibitory, mediated by the GABA_A_-dependent network mechanisms. Importantly, orexinergic input assessed by the ligand presence in the axonal terminals to the SC increased during the night time, together with the augmented sensitivity towards OXA. This higher orexinergic drive during the behaviourally active night was accompanied by lowered spontaneous neuronal activity in the SC, compared to the behaviourally quiescent day. The daily timing of these neuronal activities coincided with the timing of local rhythmic clock gene expression *ex vivo* and *in vivo*, both in the daily (under light-dark cycles) and circadian (constant dark) fashion. Additionally, we identify the OX_2_ receptor to be predominately expressed and functional in the SC, suggesting that the orexinergic pathway in the SC is arousal dependent. Together, these findings point to the broader circadian modulation at work in the SC.

## 2. RESULTS

### 2.1. Orexinergic fibres are present in the SC with a higher density of OXB-ir during the night

Despite a limited area populated by orexinergic neurons, their axons can be found throughout the whole brain. Early studies locate orexin-ir fibres at the area of the rat SC [3], but the exact spatio-temporal distribution of orexinergic innervation of distinct SC layers has not been studied before. Thus, we first performed an immunohistochemical staining against OXB on the tissue obtained from seven rats intra-ocularly injected with cholera toxin B (CtB), culled either at the beginning of light (ZT1, n=3) or dark phase (ZT13, n=4). This enabled us to observe orexinergic fibres at the retinorecipient areas of the SC, which were immunostained against CtB (Fig. 1A,B). In detail, the CtB immunostaining was found in the superficial layers, namely in the zona layer (Zo) and the full depth of superficial grey layer (SuG), contralateral to the injected eye. In all seven rats tested, OXB-ir fibres were present throughout the SC, including the superficial (Zo, SuG and optic tract layer – Op) and intermediate layers (the intermediate grey – InG and intermediate white layer – InWh).

**Figure 1.**
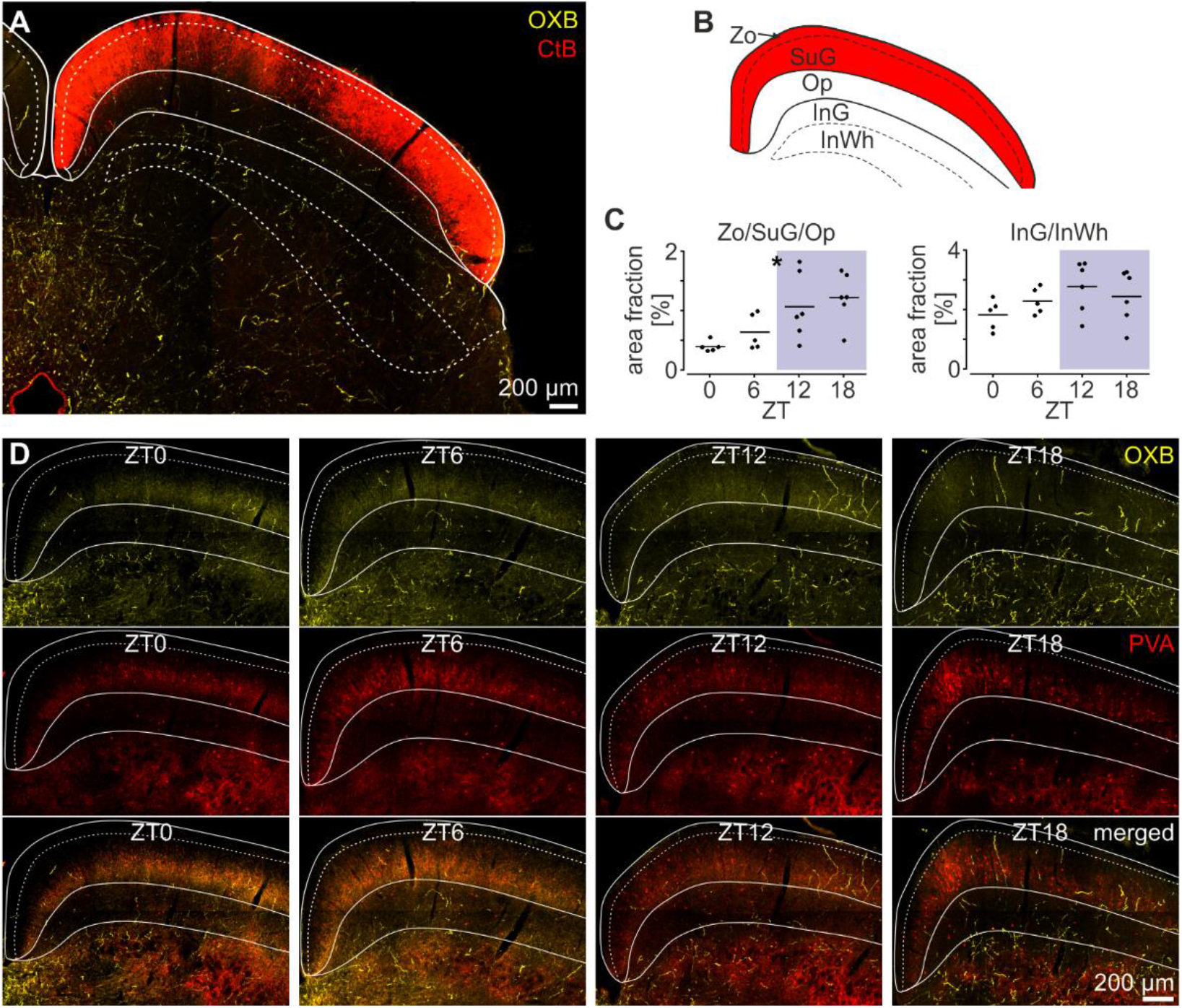
Orexin B-immunoreactive fibres are present in the SC and exhibit daily variation in its superficial layers. (**A**) Photomicrograph showing orexin B (OXB)-ir (in yellow) and retinal fibres (in red) in the midbrain after intraocular cholera toxin B (CtB) injection. The superficial and intermediate layers of the SC are outlined. (**B**) Schematic representation of the studied SC layers: Zo – the zona layer, Op – the optic tract layer, InG – the grey intermediate layer, InWh – the white intermediate layer. Red depicts the retinorecipient area of the SC. (**C**) Daily pattern of the density of OXB-ir fibres in the superficial and intermediate layers of the SC. Blue shading depicts the dark phase. (**D**) Representative confocal photomicrographs with the superficial layers outlined. OXB-ir fibres are shown in yellow, whereas red depicts parvalbumin (PVA). Merged images are shown in the bottom panel. \**p*<0.05

As the orexinergic system of the hypothalamus exhibits profound daily and circadian rhythmicity [6,41–44] and day-to-night differences in the orexin immunoreactivity were previously shown in their axons innervating the thalamus [18,19], we next tested the possibility of daily changes in the OXB-ir in the SC. To quantify the density of OXB-ir fibres in the superficial and subjacent intermediate layers of the SC, we transcardially perfused 22 rats at four time points across the day and night (ZT0 n=5, ZT6 n=5, ZT12 n=6, and ZT18 n=6). A significant variation in the density of OXB-ir fibres with a prominent night-time rise was present in the superficial layers of the SC (Zo/SuG/Op: *p*=0.0112), but this did not reach the significance threshold for the intermediate layers (InG/InWh: *p*=0.2300, one-way ANOVAs, Fig. 1C). The co-immunostaining against parvalbumin (PVA) helped us to reliably distinguish between the distinct SC layers (Fig. 1D). This dataset shows the orexinergic innervation of the entirety of the SC, including the visual superficial layers, with the latter exhibiting evident day-to-night changes in the OXB immunoreactivity.

### 2.2. The orexinergic system of the lateral hypothalamus innervates the superficial layers of the SC

The lateral hypothalamus area (LH) is established to be the primary, if not exclusive source of orexins in the brain [1–3]. Therefore, we next aimed to confirm that the LH orexinergic neurons project to the SC. Three rats were successfully injected with retrograde dyes: FluoroRed (into the left) and FluoroGreen (into the right hemisphere), targeting the superficial layers of the SC. At one week post-injection, rats were treated with colchicine 24 h before cull, to improve somatic OXB staining and minimise false negative errors. Subsequently, every third slice containing the LH was immunostained against OXB and analysed. Despite the focal and limited injection site in the medial part of the SC (Fig. 2A), we found the OXB-ir cells containing tracer granules scattered across the LH. Overall, our analysis revealed 8727 ± 188 OXB-ir cells in the hypothalamus, with 81 ± 11 filled with FluoroRed and 78 ± 5 with FluoroGreen (Fig. 2B,C). No lateralisation of OXB-ir cells filled with the retrograde dye was found in relation to the side of injection (*p*=0.0931, paired t-test, Fig. 2B). Numerous cells containing tracer granules but lacking immunoreactivity against OXB were present above the LH, thus were not further quantified. These results show that the bilateral LH is a source of the orexinergic innervation of the SC.

**Figure 2.**
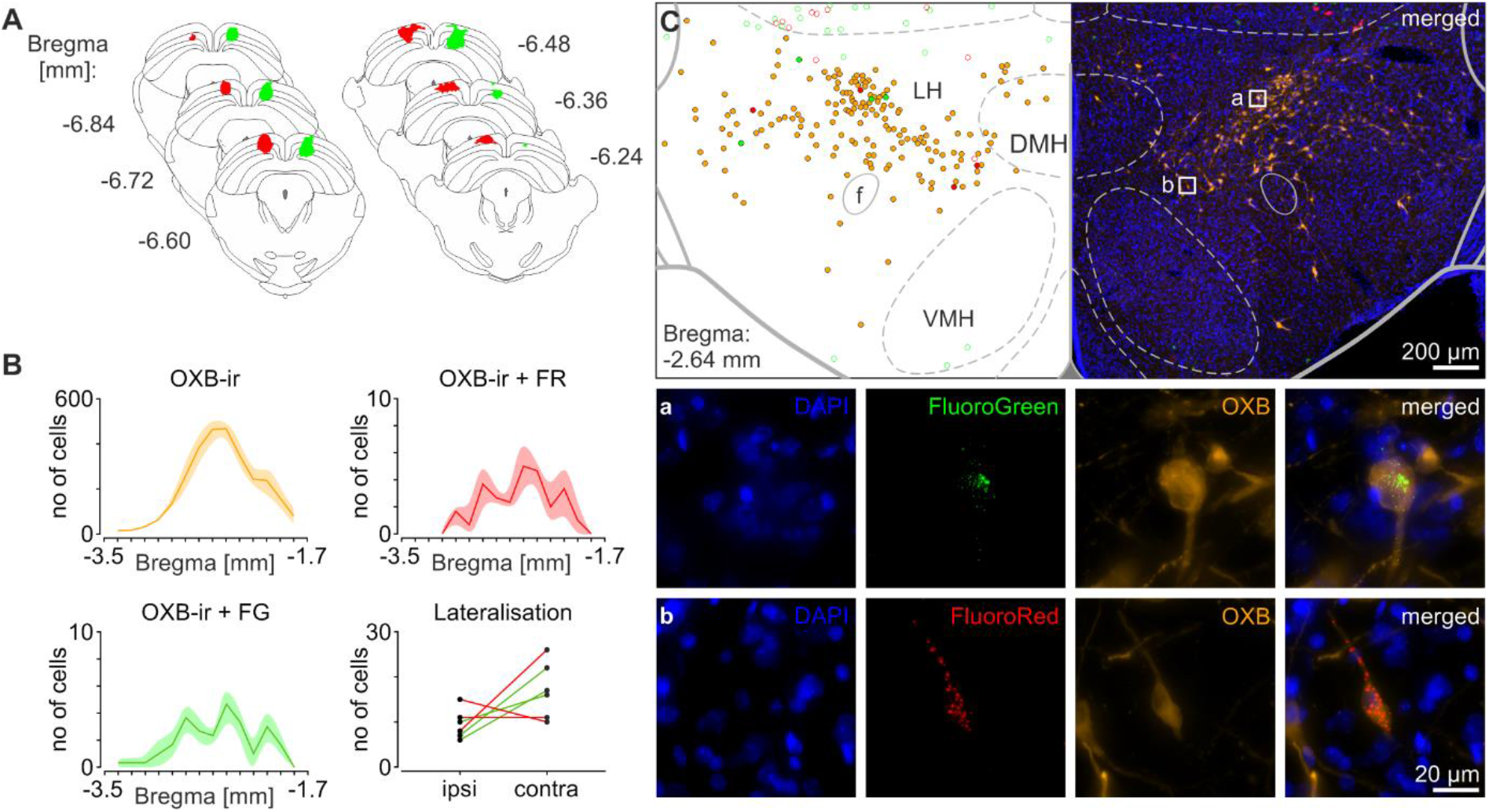
The lateral hypothalamus is the source of the orexinergic innervation of the SC. (**A**) Reconstruction of the retrograde injection site focused in the superficial layers of the SC. Red – FluoroRed (FR), green – FluoroGreen (FG). (**B**) Quantification of OXB-ir cells (in orange) in the lateral hypothalamus (LH) and these OXB-ir cells that were filled with the retrograde tracer (in red and green), in relation to Bregma (in the anteroposterior dimension). Data are presented as mean and the shading codes SEM. (**C** *left*) Reconstruction of the OXB-ir cells location (in orange) in the representative hypothalamic slice, with these co-localising FG or FR (full green and red circles, respectively). Cells filled with the tracer but OXB-negative were drawn as the empty circles (**C** *right*) Original photomicrograph. (**Ca,b**) Magnifications showing OXB-ir cells in the LH filled with FG and FR dyes. From left to right: blue – DAPI, green or red – FG or FR, orange – OXB, merged images. DMH – the dorsomedial hypothalamus, VMH – the ventromedial hypothalamus, f –fornix.

### 2.3. Orexin A modulates neuronal activity of the superficial layers of the SC in a daily fashion

The orexinergic system modulates neuronal activity of the subcortical visual system, including the lateral geniculate nucleus of the thalamus [18–22,24,45] and the SCN [46–48]. Orexins were also found to potently excite neurons in the visual cortex [49] and act at the level of the retina [50,51]. Therefore, we investigated whether the orexinergic system innervating the SC influences its spontaneous neuronal activity. To resolve this, 20 rats were culled at four time points across the daily cycle (ZT0, 6, 12, and 18; n=5 each) and the firing rates of neurons localised in the superficial layers of the SC were examined through the multi-electrode array (MEA) registrations *ex vivo*. On the whole, 1740 single units were spike-sorted from 20 recordings (20 brain slices), with 1002 neurons located in the Zo/SuG and 738 in the Op. Subsequently, their activities were evaluated following the application of OXA (200 nM), which significantly affected firing rates of the majority of neurons recorded. Both OXA-evoked activations and inhibitions of single unit activity (SUA) were found in the Zo/SuG and Op (Fig. 3A,C,D), with the first layer being significantly more responsive to the treatment than the latter (*p*<0.0001, Chi^2^ test, Fig. 3E,G). However, when SUA was averaged, the application of OXA evoked the overall activation of the SC (Fig. 3B). This implies that on the level of a whole structure, these OXA-evoked excitations prevail inhibitions.

**Figure 3.**
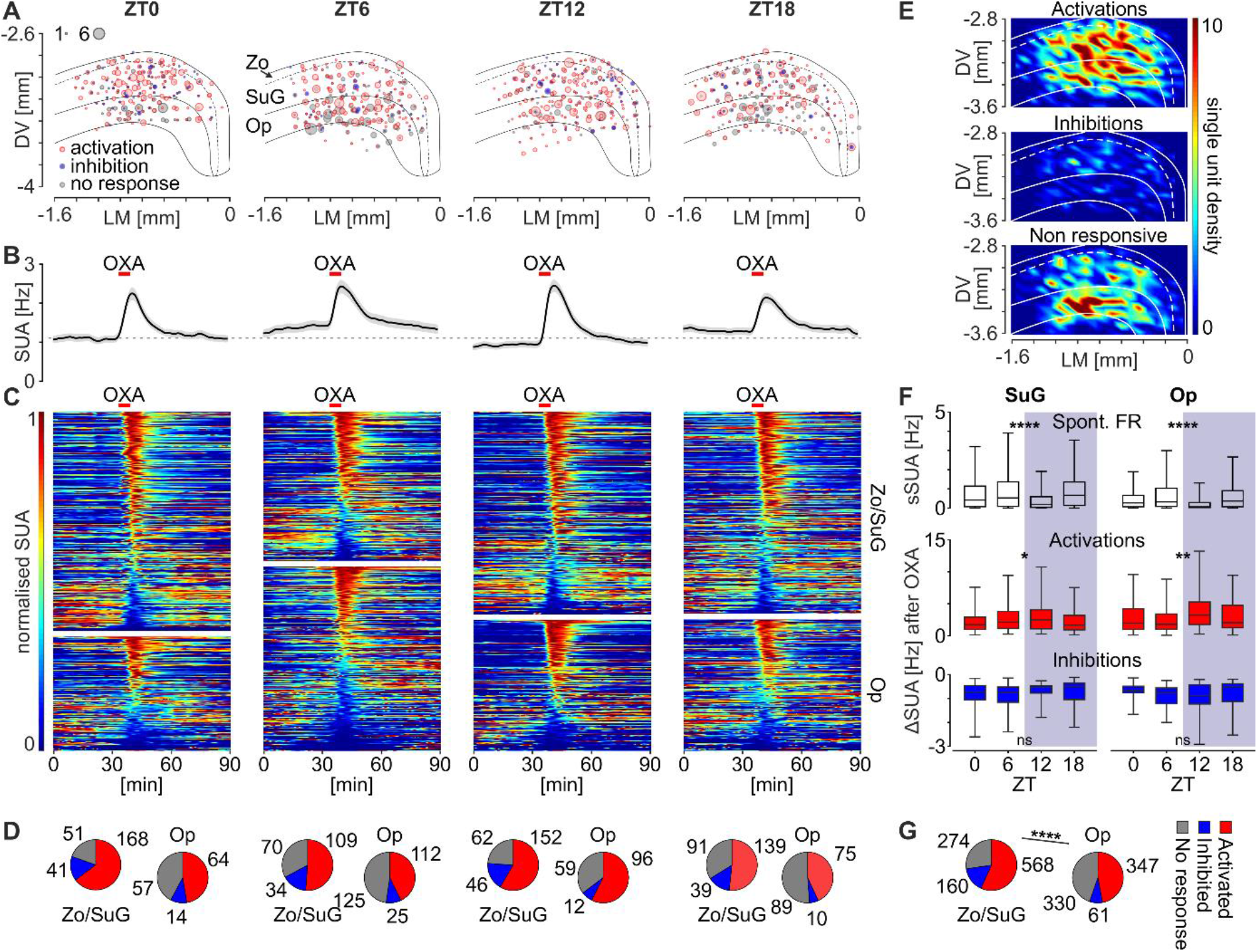
Orexin A modulates the neuronal activity of the majority of SC neurons populating its superficial layers, depending on the time of a day. (**A**) Spatial bubble density plots showing single units activated (in red), inhibited (in blue) or not affected (in grey) by the application of OXA (200 nM) at each ZT, with the superficial layers of the SC outlined. The size of a bubble codes the number of units pulled from a small subregion. Zo – the zona layer; SuG – the superficial grey layer; Op – the optic tract layer; DV – dorsolateral; LM – lateromedial. (**B**) Mean effects of OXA in each daily time point, shown as single unit activity (SUA) histograms (bin: 30 s). Single units were pulled from five recordings per time point and sorted from top to bottom based on the relative SUA during the first response to OXA. Data presented as mean ± SEM. Red bars mark the time of OXA application. (**C**) Heatmaps showing the normalised SUA for each recorded neuron in the Zo/SuG and Op (bin: 30 s). Warm colours code the relatively high, whereas cold – low firing rates. (**D**) Pie charts displaying the proportion of OXA-evoked activations (in red), inhibitions (in blue) and these cells, which did not respond to the treatment (in grey). (**E**) Spatial density heatmaps for all single units recorded in four time points, with the borders of superficial SC layers outlined. Warm colours code high density of units that were activated, inhibited or not affected by OXA, respectively (*from top to bottom*). (**F**) Box plots showing daily changes in the spontaneous firing rate and the amplitude of OXA-evoked activations and inhibitions. Whiskers were set to encompass all data points. (**G**) Pie charts summarising the responsiveness of the superficial layers of the SC to OXA, regardless of the time of a day. \**p*<0.05, \*\**p*<0.01, \*\*\*\**p*<0.0001.

Both the spontaneous firing rate and the amplitude of response to OXA were subject to a daily change, with distinct temporal patterns. Neurons localised in both the Zo/SuG and Op exhibited a pronounced drop in spontaneous SUA at early night, compared to the day, which rebounded at ZT18 (*p*<0.0001, Kruskal-Wallis tests, Fig. 3F). The amplitude of OXA-evoked activations exhibited daily changes in an opposite phase, reaching its peak at ZT12 (Zo/SuG: *p*=0.0157, Op: *p*=0.0029, Kruskal-Wallis tests, Fig. 3F). No day-to-night changes in the amplitude of OXA-evoked inhibitions were noted (Zo/SuG: *p*=0.1124, Op: *p*=0.5011, Kruskal-Wallis tests, Fig. 3F). These data demonstrate a prominent modulatory action of OXA on the neuronal activity of the superficial layers of the SC. This modulatory action occurs with daily variation such that the spontaneous activity of SC neurons and their responsiveness to OXA are oppositely timed around the daily cycle.

### 2.4. Orexin A-evoked inhibitions of the SC neuronal activity depend on GABAergic mechanisms

Direct action of orexins upon a variety of neuronal subpopulations in the central nervous system is predominately excitatory [4], with the postsynaptic hyperpolarisation of clock cells in the SCN by OXA remaining an exception [46]. The SC contains multiple subpopulations of GABAergic neurons, and the SuG in particular is populated by GABAergic interneurons that do not project outside the SC, but instead provide horizontal inhibitory tone inside this brain structure [52,53]. To evaluate if OXA-evoked inhibitions of SUA in the SC are mediated by these GABAergic connections, we performed subsequent applications of OXA (200 nM) in control conditions and in the presence of GABA_A_ receptor antagonist, bicuculline (20 μM) on three brain slices recorded on the MEA *ex vivo*. Pre-treatment with bicuculline temporarily disinhibited neuronal activity in the SC, what proves the preservation of GABAergic network in the studied brain slices (Fig. 4A,B). In the presence of GABA_A_ receptor blockage, OXA failed to evoke inhibitions in SUA, or the amplitude of these inhibitions was significantly reduced (*p*<0.0001, Friedman test followed by Dunn’s multiple comparison, Fig. 4C). On the contrary, the amplitude of OXA-evoked activations was unaffected by the antagonist (*p*=0.7124). Results of this experiment provide evidence on the GABAergic network-dependent nature of this OXA-evoked inhibition of the SC neuronal activity.

**Figure 4.**
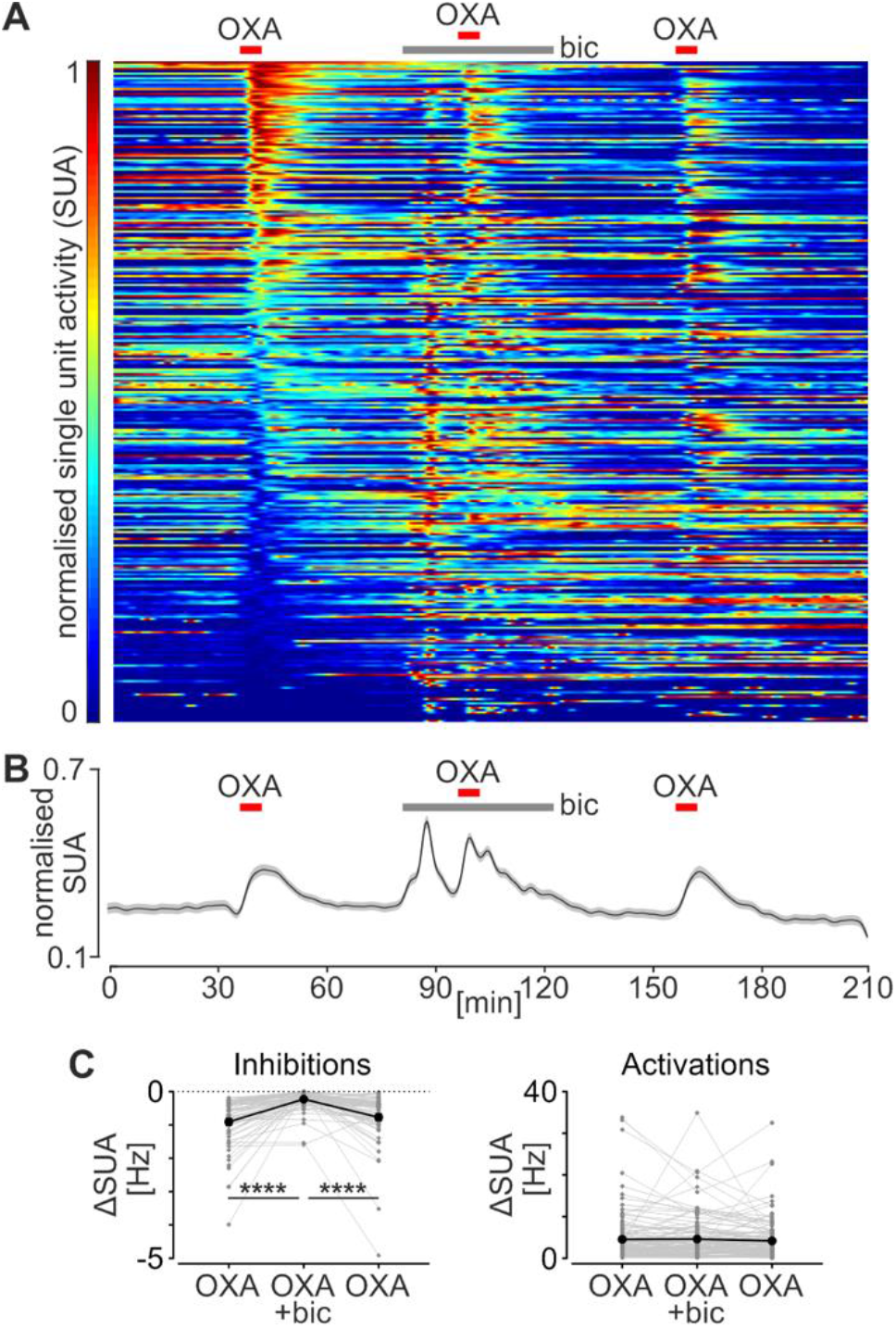
Orexin A-evoked inhibitions are mediated by the GABAergic network in the SC. (**A**) A heatmap displaying the normalised SUA (bin: 30 s) during a triple OXA (200 nM, red bar) application. Grey bar points the time of a tonic bicuculline (bic, 20 μM) application. Units were sorted from the top to bottom according to the relative activity during the first response to OXA. (**B**) Average plot (bin: 30 s) of the normalised SUA. Data were drawn as mean ± SEM. (**C**) Amplitude of the responses, shown separately for inhibitions and activations. \*\*\*\**p*<0.0001.

### 2.5. Orexin A modulates neuronal activity in the SC by the activation of OX_2_ receptor

Orexins act on two metabotropic G-protein coupled receptors: OX_1_ and OX_2_ receptor [1,2,54], with OXA binding to these two with similar affinity. As the expression of orexin receptors varies across brain structures, we next set up to establish the predominant receptor functional in the SC. First, we performed triple OXA (200 nM) applications on five SC slices: in control conditions, in the presence of a specific OX_2_ receptor antagonist TCS-OX2-29 (10 μM), and after its washout. Both OXA-evoked activations (n=212) and inhibitions (n=29) were eliminated by TCS-OX2-29, and restored after the antagonist washout (*p*<0.0001, Friedman tests followed by Dunn’s multiple comparisons, Fig. 5A). Second, we tested whether the OX_1_ receptor antagonist OXA17-33 (1 μM) modulates the amplitude of OXA-evoked response of SC neurons. The application of the antagonist upon two SC slices failed to reduce both activations (n=48, *p*=0.0742) and inhibitions in the response to OXA (n=14, *p*>0.9999, Dunn’s multiple comparisons, Fig. 5B). Finally, we tested if repetitive OXA applications cause orexin receptors to desensitise, by applying OXA three times without any drugs added on one SC slice. The amplitude of OXA-evoked activations (n=44, *p*=0.3284) and inhibitions (n=13, *p*=0.3679, Friedman tests, Fig. 5C) did not significantly change over time. These data provide evidence for the functional expression of OX_2_ receptors in the superficial layers of the SC.

**Figure 5.**
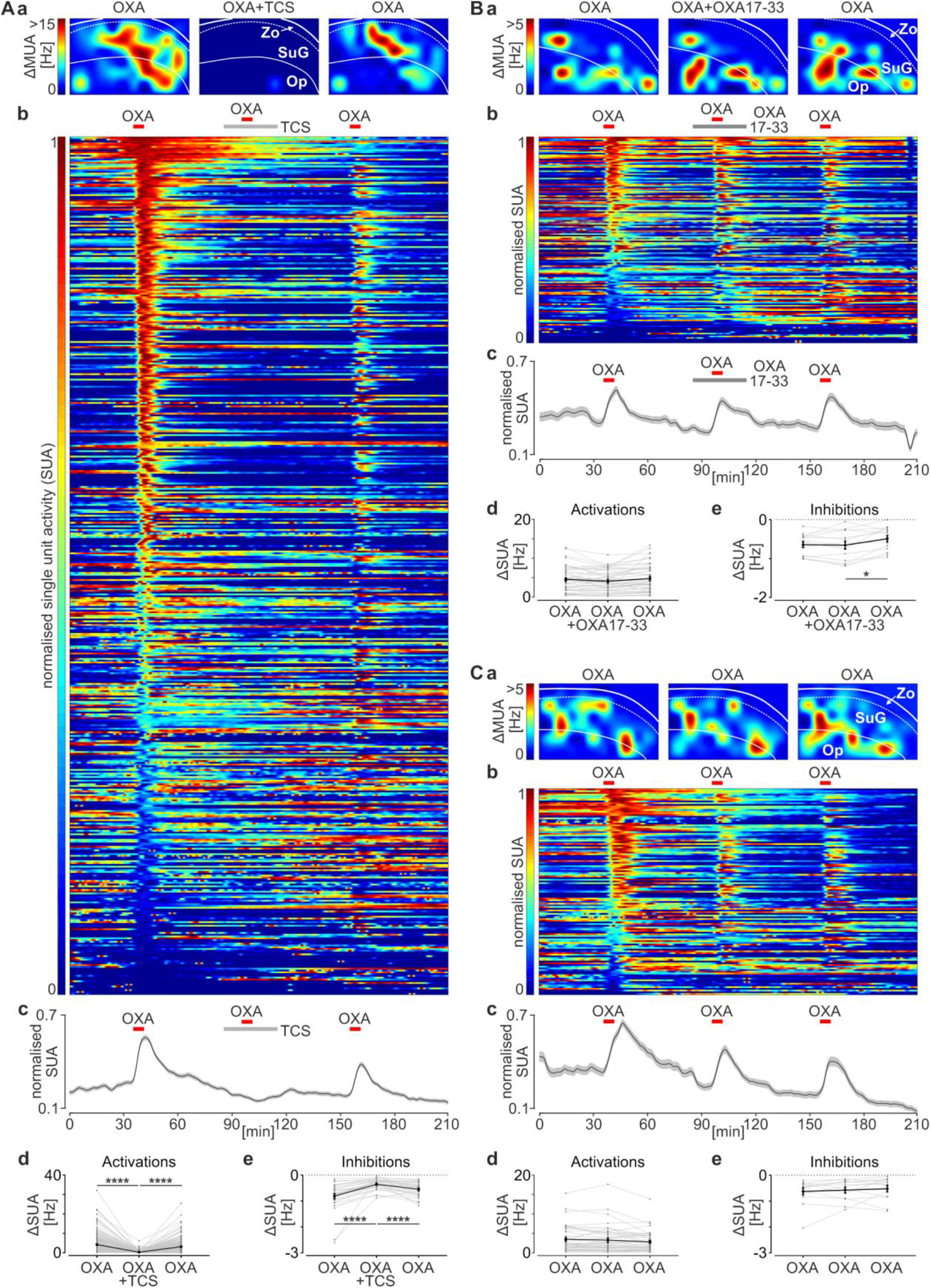
OX_2_ receptor is predominant in the superficial layers of the SC. (**A**) The effects of OX_2_ receptor antagonist TCS-OX2-29 (TCS, 20 μM) on the response amplitude to OXA (200 nM). (**B**) OX_1_ receptor antagonist OXA17-33 (1 μM) failed to reduce the effects of OXA on the SUA. (**C**) Control, triple applications of OXA. (**a**) Spatial heatmaps showing the change in multi-unit activity (MUA) after OXA application. The superficial layers of the SC were outlined according to the localisation of the multi-electrode array. (**b**) Temporal heatmaps displaying normalised SUA throughout the recording (bin: 30 s). Units were sorted from top to bottom according to relative activity during the first response to OXA. (**c**) Average plots of the normalised SUA. Data were drawn as mean ± SEM. (**d,e**) Statistical analyses of the amplitudes of OXA-evoked activations (*d*) and inhibitions (*e*): before, during and after the treatment with the antagonist, or during the triple control application of OXA. \**p*<0.05, \*\*\*\**p*<0.0001.

### 2.6. The superficial layers of the SC possess molecular clock mechanisms which are sustained in constant darkness

Neurons of the central clock but also at an emerging number of extra-SCN brain centres are being shown to express clock genes with 24 h period, to organise their circadian physiology. This includes daily rhythms in the firing rate or in the expression of different receptors, leading to day-to-night changes in the responsiveness to a variety of neurotransmitters and neuromodulators [9,10,12,15,16]. Therefore, we next aimed to establish whether the superficial layers of the SC express the molecular clock *in vivo*, which enables SC neurons to regulate their circadian physiology, including their neuronal activity and responsiveness to orexins. First, 24 rats were maintained under the standard 12:12 light-dark cycle (LD) and culled in four daily time points: ZT0, 6, 12, and 18 (n=6 each). Both *Per1* and *Bmal1* expression in the SC varied significantly over these time points (*Per1*: *p*=0.0005; *Bmal1*: *p*=0.0295, one-way ANOVA, Fig. 6A). Daily pattern of expression for these two clock genes remained close to the antiphasic relationship, with *Per1* peaking at ZT12 and *Bmal1* at ZT0. Next, another cohort of 24 rats was moved from these standard LD conditions to the constant darkness (DD) for 48 h before being culled in four circadian time points: circadian time (CT)0, 6, 12, and 18 (n=6 each). Under these conditions, the expression of *Per1* and *Bmal1* remained significantly changing across 24 h (*Per1*: *p*=0.0008; *Bmal1*: *p*<0.0001, one-way ANOVA), with a peak in *Per1* coinciding with a nadir in *Bmal1* expression level. To gain more insights into the circadian expression of clock genes in the SC, we additionally measured the expression level of *Rev-erbα* and *Clock*, which also significantly varied across CT (*Rev-erbα*: *p*<0.0001; *Clock*: *p*<0.0001, one-way ANOVA, Fig. 6B). Similarly to *Bmal1*, the expression of both clock genes was the lowest at CT12, the time of *Per1* acrophase. Thus, the rhythmic expression of clock genes in the SC follows a daily and circadian pattern *in vivo*.

**Figure 6.**
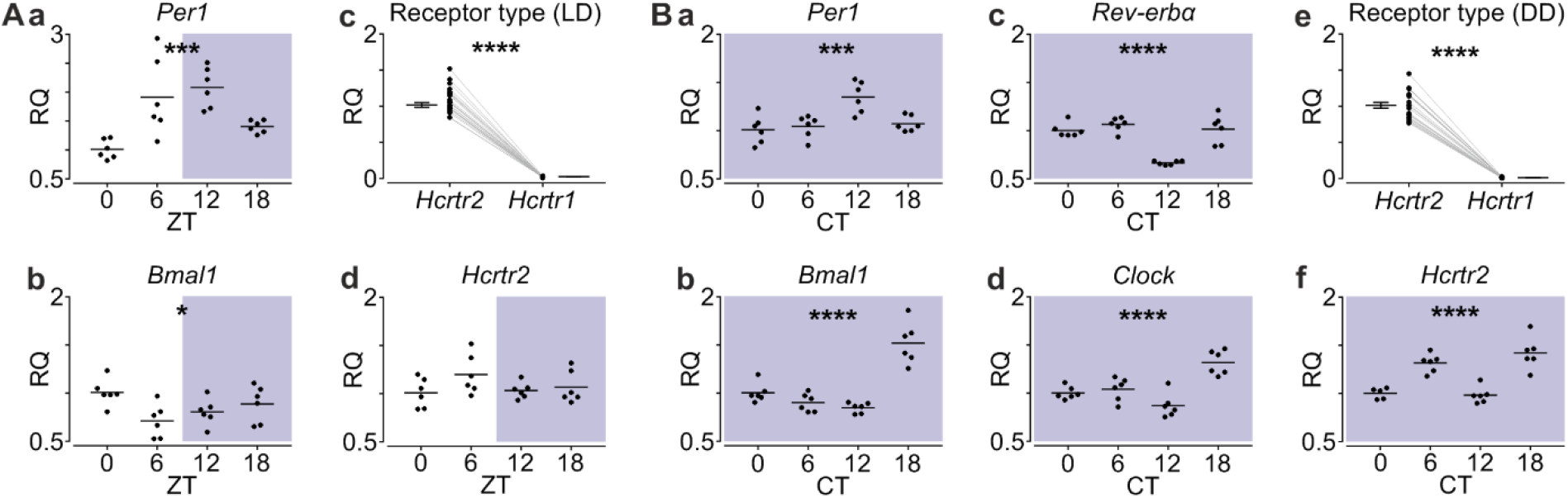
The superficial layers of the rat SC express core clock genes in the daily and circadian fashion. (**Aa,b**) Expression of core clock genes *Per1* and *Bmal1* under standard 12:12 LD cycle. (**Ac**) The quantitative comparison of orexin receptor genes expression. (Ad) Expression of the prevalent *Hcrtr2* under LD. (**Ba-d**) Expression of clock genes *Per1, Bmal1, Rev-erbα* and *Clock* under DD. (**Bc**) The comparison of *Hcrtr1* and *Hcrtr2* expression under DD. (**Bf**) Expression pattern of the dominant *Hcrtr2* under DD. Blue shading codes the darkness. ZT – Zeitgeber time, CT – circadian time, RQ – relative gene expression; \**p*<0.05, \*\*\**p*<0.001, \*\*\*\**p*<0.0001.

Under both LD and DD conditions, we additionally measured the level of orexin receptor gene expression. In these SC samples, the expression of OX_2_ receptor gene (*Hcrtr2*) was approximately 50 to 100-fold higher than that of OX_1_ receptor gene (*Hcrtr1*) (*p*<0.0001, paired t-tests, Fig. 6A,B). Moreover, the expression of the prevalent *Hcrtr2* varied significantly under DD (*p*<0.0001, Fig. 6B) but not LD conditions (*p*=0.1939, one-way ANOVAs, Fig. 6A). This dataset is in keeping with our electrophysiological results pinpointing the functional OX_2_ receptors express dominantly in the SC.

### 2.7. Clock cells in the superficial layers of the SC express OX_2_ receptors

The SC stands a powerful computational network of different neuronal subpopulations creating organised connections within this structure [53]. Thus, we next aimed to unravel if OX_2_ receptors are expressed by the same SC neurons which organise their temporal physiology due to clock gene expression. To address this, we utilised RNAscope technology – the fluorescent *in situ* hybridisation, on SC slices obtained from six rats. For transcript detection we used three different probes: against *Per2* (a selected clock gene), *Hcrtr2* (OX_2_ receptor gene) and *Penk* (the proenkephalin gene, used for delineation of distinct SC layers). Clusters of *Hcrtr2* and *Per2* around DAPI-stained nuclei were present in the whole extent of the superficial layers of the SC, particularly in the SuG and Op. Further visual inspection revealed high co-localisation of *Per2* and *Hcrtr2* around the same nuclei of SC cells (Fig. 7). Altogether, the RNAscope data strongly supports that clock cells in the visual SC express OX_2_ receptors.

**Figure 7.**
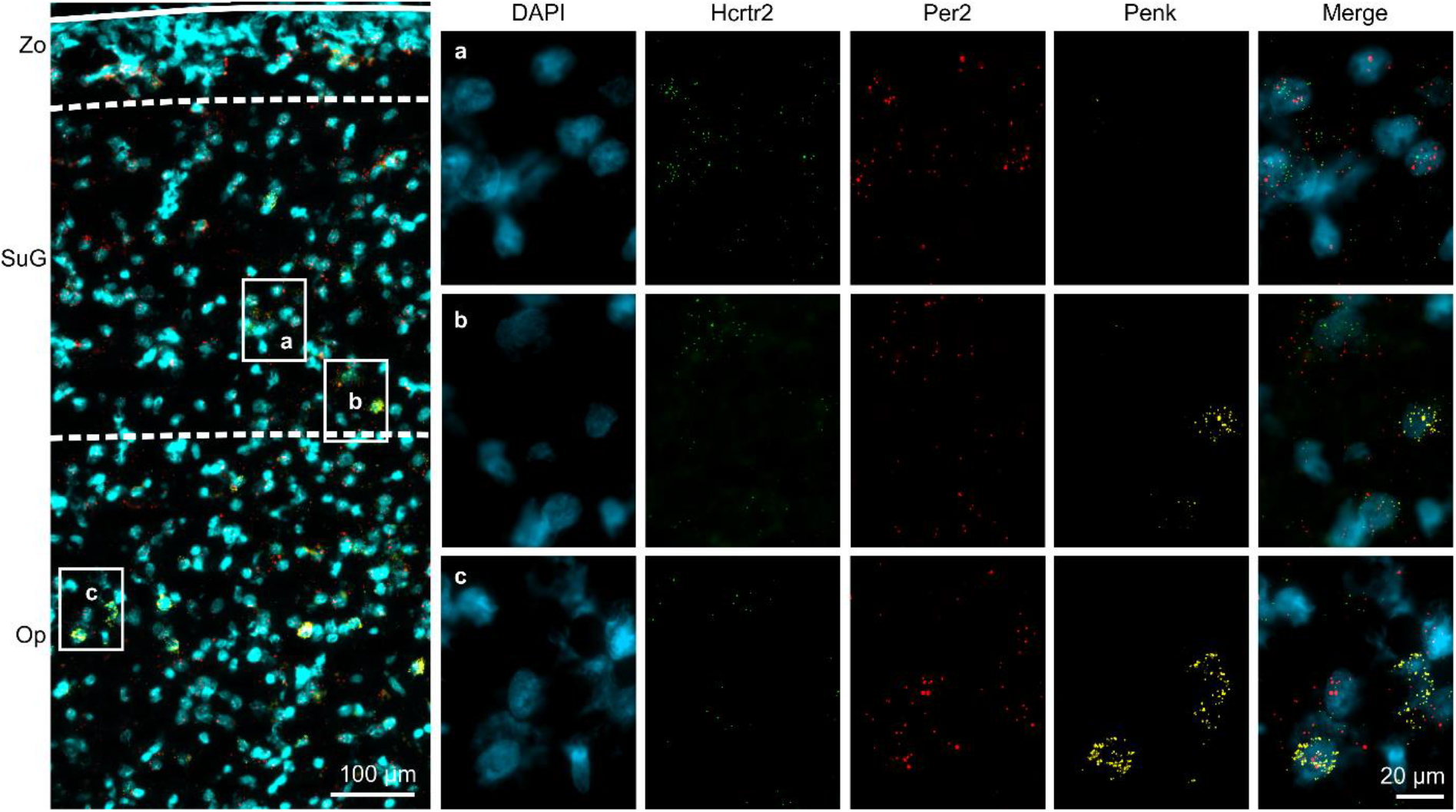
Co-expression of OX_2_ receptor gene *Hcrtr2* with a core clock gene *Per2* in the superficial layers of the SC. (*left panel*) Low magnification photomicrography with superficial layers of the SC outlined: Zo – the zona layer; SuG – the superficial grey layer; Op – the optic tract layer. (*right panel*) High magnification of two inserts from the SuG (*a,b*) and one from the Op (*c*). Nuclei counterstained with DAPI are presented in cyan, *Hcrtr2* in green, *Per2* in red, and *Penk* (used as a marker for SC layers distinction) in yellow. The merged signal shows high co-localisation of *Hcrtr2* and *Per2*, particularly in the SuG.

### 2.8. Intrinsic circadian oscillations in core clock gene expression in the mouse SC

Finally, we aimed to resolve whether rhythmic clock gene expression in the SC *in vivo* arises from local, intrinsic clock mechanisms in this structure, or is a consequence of the phase communication from another brain clock [55]. Therefore, we used a PERIOD2∷LUCIFERASE mouse model, where the *Per2* expression is monitored through PER2∷LUC bioluminescence [56]. Under isolated culture condition, the SC continued to maintain the molecular circadian clock for up to six days (Fig. 8). Interestingly, the posterior SC in our preparation, corresponding to the area used in our electrophysiological protocols carried out on rats, was seen to maintain a higher amplitude bioluminescence oscillation than the anterior SC. These results indicate that SC harbours intrinsic circadian clock mechanisms independent from a daily input from other brain areas.

**Figure 8.**
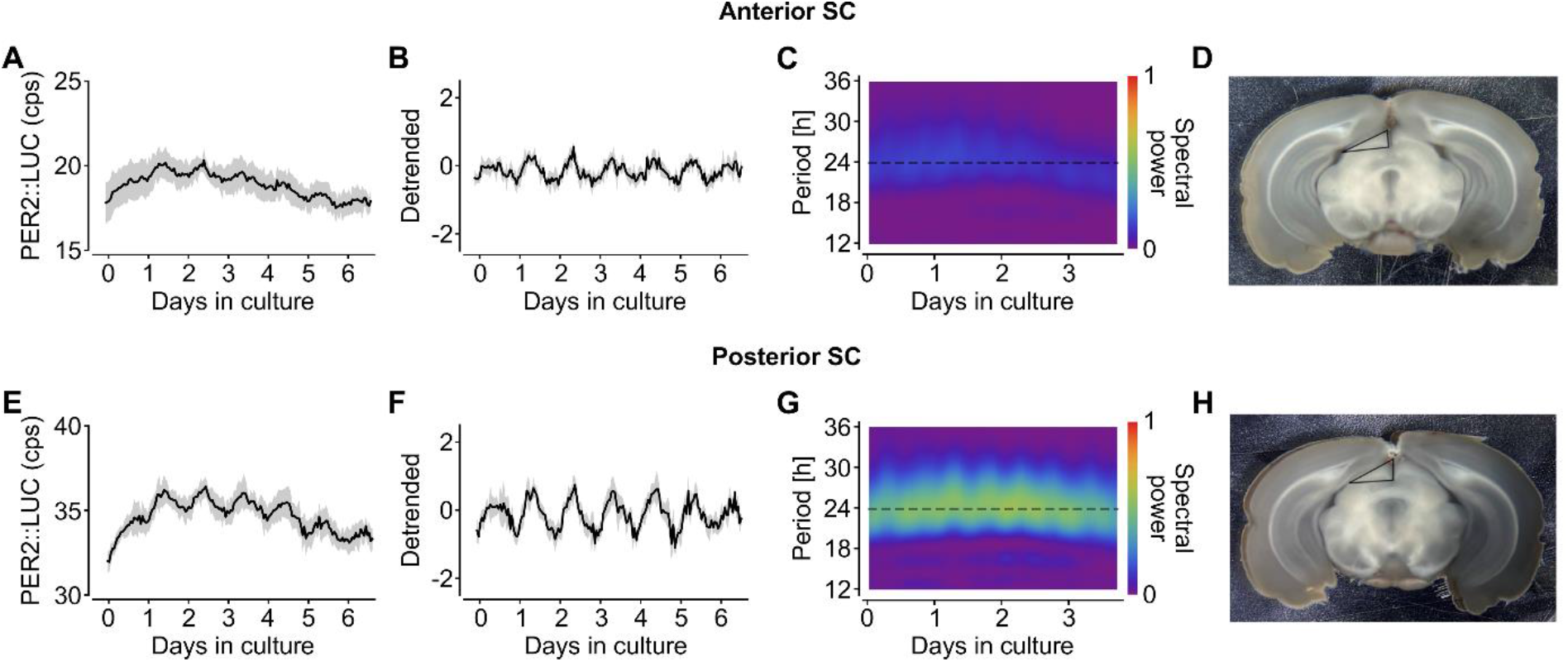
Anterior and posterior coronal sections of the mouse SC maintain circadian clock *ex vivo*. **(A)** PER2∷LUC expression maintains circadian oscillation in the explant culture of the anterior SC. The solid line represents an ensemble average (n=4 animals) and the shade indicate the standard deviation. (B) Detrended PER2∷LUC signal. (C) The spectrogram through the sliding FFT reveals a circadian spectral power around 24-h period in the anterior SC (ensemble average over spectrograms, n=4 animals). (D) The mouse brain slice containing the anterior SC. The triangled area of the superficial layers of the SC was sampled for explant culture. (E) Circadian PER2∷LUC oscillation is also observed in the posterior SC explant with a higher amplitude than the anterior SC. (F) Detrended bioluminescence. (G) A strong circadian spectral power in the posterior SC continues during culture, with stronger power than that of the anterior SC. To enable comparison of spectral power between the two SC explants, the colour scales in (*C*) and (*G*) are set to be identical. (H) The mouse brain slice containing the posterior SC, indicated by the triangled area explanted for culture.

## 3. DISCUSSION

In this study, we reveal modulatory actions of orexinergic system on the superficial layers of the SC and observe temporal changes in the SC physiology that occur in the daily timescale. The orexinergic innervation originating most likely from the lateral hypothalamus increases past the day-night transition point (ZT12), which coincides with the timing of higher activation of SC neurons by OXA. We also found spontaneous neuronal activity in the SC to vary across the daily cycle, with lowest firing rates at early night corresponding to an increased sensitivity to OXA. These temporal events are highly likely to be coordinated by local circadian clocks. Indeed, we verified the presence of the molecular circadian clock in the SC, exhibiting daily and circadian rhythmicity both *in vivo* and *ex vivo*.

We first show the presence of OXB-ir fibres at the whole extent of SC layers, including the retinorecipient ones, confirming early anatomical studies [3,57]. The number of immunoreactive axons could be classified as sparse (in the superficial layers) to moderate (in the intermediate and deep layers). However, the low number of fibres does not necessarily indicate orexins to act on a limited number of SC cells; the volume rather than synaptic transmission is widely reported for peptides including orexins [58]. The robust release of these neuropeptides into the SC is supported by the report showing the highest concentration of OXA in the SC tissue homogenate amongst all the brain areas studied outside the hypothalamus [59]. Moreover, using retrograde neuronal tract tracing methods, we recognise the LH as a source of the orexinergic innervation of the SC. The moderate number of cells scattered throughout the extent of the hypothalamus corresponds both to the limited site of tracer injection and the density of orexinergic axons found in superficial layers of the SC.

In keeping with the observation that the orexin release to the SC is relatively high despite a moderate number of fibres [59], we found that OXA is a potent modulator of the neuronal activity of most neurons localised in the superficial layers of the SC. This was true for both the Op and Zo/SuG, with the latter significantly more affected by OXA. Of note, these effects were not only excitatory, but 12.7% of the recorded single units were notably inhibited by the application of OXA. Since a direct orexin-evoked hyperpolarisation is very rare and up to date characterised for the SCN only [46,60], we tested and confirmed that inhibitions in the SC SUA were abolished by the blockage of GABA_A_ receptors. This revealed that even in the slice preparation *ex vivo*, an intact GABAergic network functions in the SC [52,53] to propagate the orexinergic signal to cells further into the network, which do not express orexin receptors on their own.

In the central nervous system, the expression of distinct orexin receptors shows a largely compartmentalised pattern amongst brain structures, which implies divergent physiological roles played by these two receptor types. Our molecular, *in situ* and electrophysiological evidence supports the expression of orexin receptors in the superficial layers of the SC, with the pronounced (up to 100-fold) predominance of the OX_2_ over the OX_1_ receptor. OX_2_ receptors were previously reported to be expressed in the SC with the use of *in situ* hybridisation and immunohistochemistry [61,62]. Additionally, no OX_1_ receptor-positive cells were detected in the SC of OX_1_R-eGFP mice [63]. The predominant action of OXA via the OX_2_ receptor is shared by other retinorecipient subcortical brain centres as the dorsal and ventral nuclei of the lateral geniculate complex [18–20]. Functionally, the OX_2_ receptor is deemed responsible for mediating the arousal-related actions of orexins, whereas the expression of OX_1_ receptor was linked to their impact on food intake [64]. Therefore, the orexin action within the SC can be linked to behavioural arousal.

A plethora of physiological processes including those under control of the orexinergic system are subject to daily/circadian modulation. The master clock localised in the SCN orchestrates most of the circadian rhythms in the brain and body, due to the cycling expression of clock genes [65–67]. However, accumulating evidence describes multiple extra-SCN brain oscillators, which autonomously or semi-autonomously express clock genes to coordinate their processes in a day-to-night changing manner [9,11]. In our study, we describe for the first time the daily (under LD) and circadian (under DD conditions) expression of clock genes in the SC, such as core *Per1*, *Bmal1* or *Clock* and accessory *Rev-Erbα*. Their phasic interrelations follow the transcriptional logic of the SCN or other extra-SCN clocks: the phasing of *Rev-Erbα* is close to *Bmal1* and *Clock*, as their transcription is initiated by Rev-ErbA/ROR response element, while their expression stands in an antiphase to that of *Per1* expression governed by E-box [10]. This indicates that the transcriptional-translational feedback loop is at work in the SC. However, clock gene expression may be organised *in vivo* by a rhythmic input and not necessarily be synchronised enough by local intrinsic mechanisms to produce a well-structured phasic output at the level of a whole brain area; a mismatch seen before for different timekeeping brain areas [55,56]. Thus, evidence for the existence of local circadian clock in the SC was strengthened by real-time PER2∷LUC expression monitoring of the SC explant culture, where the bioluminescence reporting *Per2* expression was rhythmic up to six days. Thus, the SC molecular clock operating independently of remote brain clocks may result in the autonomous circadian modulation of electrophysiological activities in the SC brain slice.

Our MEA recordings unravelled the daily variation in the spontaneous neuronal activity in the superficial layers of the SC, exhibiting a pronounced decrease at the beginning of dark phase, compared to the high levels of SUA throughout the day. It is unlikely for this change to be caused by reduced excitatory retinal input in the darkness, as these rats were culled exactly at ZT12, thereby not being affected by dark conditions for longer than a few minutes. Moreover, the SC SUA rebounded in the middle of the night, which suggests this daily variation comes endogenously, and is not a simple fingerprint of the environmental light conditions before the cull. The daily fluctuation in the SC firing rate can be thus attributed to the local clock gene expression *in situ*, which may reorganise *in vivo* when it receives rhythmic inputs from the central clock.

The orexinergic system of the hypothalamus also expresses daily and circadian changes, as reflected in the rhythmic neuronal activities and responsiveness to environmental light conditions [6]. The arousal-promoting orexinergic neurons are more active during the behaviourally active phase, which is paralleled by the increase in OXB-ir in the SC, shown by our study. Additionally, at the beginning of the night, when the orexinergic input to the SC significantly rises compared to the day, neurons in its superficial layers exhibit the peak in sensitivity to OXA. The daily/circadian change in the response to a peptide may stem from an intrinsic regulation of the receptor expression or alternatively, it may be a result of indirect neurophysiological changes, including the variation in membrane resistance or resting potential [15,47,68]. Our study shows the same SC neurons to co-express OX_2_ receptors and clock gene *Per2*. However, the variation in the transcriptional expression level of *Hcrtr2* did not exhibit a standard daily pattern (rather a bimodal distribution) and this variation was only significant for DD conditions. Therefore, the daily change in responsiveness to OXA in the SC should not be attributed to changes in the orexin receptor genes expression alone.

From the physiological perspective, the subcortical visual system experiences extreme day-to-night changes in its main excitatory input, which originates from the retina. Therefore, not only the detection of light, but also the anticipation of the transition to light phase, creates a huge evolutionary advantage for the animal. Our results show that the early-night increase in both the sensitivity to orexins and the orexinergic input is temporally aligned to a drop in neuronal activity in the retinorecipient SC layers expressing clock genes. We theorise that these enable the nocturnal animal to predict and detect limited light intensity in the environment during this acute behavioural arousal phase of the night. As the main function of the SC is the coordination of movements towards the visual stimulus [26,30], the facilitation of a sparse visual information under low ambient lighting must be crucial.

In both the basic research and clinical studies, the level of activity of the superficial layers of the SC has been negatively correlated with the expression of visual hallucinations [30–32,69]. Interestingly, the same symptom is shared by narcoleptic patients, whose orexinergic system malfunction is most often attributed to OX_2_ receptor loss, and by patients undergoing the insomnia treatment with DORAs [33,34,36]. As here we provide evidence for the predominately excitatory action of OXA upon the visual SC layers via the OX_2_ receptor, we speculate that the reduction of this excitatory drive in narcolepsy and during the pharmacological treatment with non-specifically acting DORAs may attribute to the expression of visual hallucination in these patients.

In conclusion, the results of our study describe and characterise visual layers of the SC to undergo daily modulation by the orexinergic system of the hypothalamus and identifies that the SC neuronal activities are shaped by both the external and intrinsic timekeeping mechanisms. Moreover, we provide physiological and pharmacological perspective for the orexinergic excitation of SC neurons via the OX_2_ receptor to support arousal during the behaviourally active dark phase, when the detection of sparse ambient light is critically needed.

## 4. MATERIALS AND METHODS

### 4.1. Animals and ethical approval

This study was performed on 106 adult male Sprague Dawley (10-12 weeks old) rats kept under a standard 12:12 light-dark cycle, unless stated otherwise, at 23 ± 2°C and 67 ± 5% relative humidity. Animals were fed *ad libitum* and had free access to water. Rats were bred in-house by Animal Facility of the Institute of Zoology and Biomedical Research at the Jagiellonian University in Krakow. All procedures were approved by the Local Ethics Committee and were performed in accordance with Polish and the European Council Directive (86/609/EEC). Procedures performed in darkness (including culls) were carried out with the use of night vision infrared goggles (Pulsar, Vilnius, Lithuania).

PERIOD2∷LUCIFERASE (PER2∷LUC) mice (RRID: IMSR_JAX:006852) were kept under a 12:12 light-dark cycle at 21 ± 2°C and 59 ± 4% relative humidity in Taipei Medical University Laboratory Animal Centre. The animals and protocols were reviewed and approved by the Institutional Animal Care and Use Committee of Taipei Medical University (IACUC Approval No: LAC-2019-0118). All experiments were designed to minimise the number of animals used and their sufferings.

### 4.2. Intraocular injections of cholera toxin B subunit (CtB)

#### 4.2.1. Surgery

Surgery for the intraocular injections of the CtB were performed as described previously [18]. In brief, seven rats were deeply anaesthetised with isoflurane (3% v/v air mixture; Baxter, Deerfield, IL, USA) and placed in the gas free-flow mask throughout the procedure. Intraocular injections of 2 μl of CtB (0.5% m/v in saline; Sigma, Darmstadt, Germany) were performed into the vitreous chamber of one eye. Animals were intramuscularly injected with Torbugesic (0.2 mg kg^−1^ body weight; Zoetis, Parsippany, NJ, USA) and Tolfedine 4% (4 mg kg^−1^ body weight; Biowet, Puławy, Poland) and returned to the Animal Facility for three days with free access to drinking water containing the antibiotic Sul-Tridin 24% (1:300; sulphadiazine 200 mg ml^−1^ + trimethoprim 40 mg ml^−1^; ScanVet, Poland).

#### 4.2.2. Tissue preparation and immunostaining

Next, rats were deeply anaesthetised with sodium pentobarbital (2 ml kg^−1^ body weight; Biowet) and transcardially perfused with 4% paraformaldehyde (PFA) solution in 0.1 M phosphate-buffered saline (PBS) at the beginning of the day (Zeitgeber time, ZT1, n=3) or night (ZT13, n=4). Subsequently, the brains removed from the skulls were post-fixed in the same solution overnight. Blocks of tissue containing midbrain were cut on the vibroslicer (VT1000S; Leica Microsystems, Wetzlar, Germany) on 40 μm thick coronal slices. Slices were rinsed in PBS and permeabilised with 0.5% Triton-X100 (Sigma) solution in PBS, additionally containing normal donkey serum (NDS; Abcam, Cambridge, UK) at room temperature for 35 min. Then, slices were transferred to a PBS solution composed of 0.1% Triton-X100, 0.5% NDS and primary antisera: goat anti-OXB (1:500; Santa Cruz Biotechnology, Santa Cruz, CA, USA) and mouse anti-CtB (1:250, Abcam). After 24 h of incubation in 4°C, sections were rinsed in fresh PBS and kept in the secondary antibodies solution (donkey anti-goat Alexa Fluor 467 and donkey anti-mouse Cy3, 1:300; Jackson ImmunoResearch, West Grove, PA, USA) overnight at 4°C. Last, slices were washed off in PBS, mounted on glass slides and coverslipped with Fluoroshield with DAPI (Sigma). Images were collected using the A1Si confocal laser scanning system (Nikon, Tokyo, Japan) built on an inverted Ti-E microscope (Nikon).

### 4.3. Daily immunohistochemical studies

#### 4.3.1. Tissue preparation and immunostaining

A cohort of 22 Sprague Dawley rats were anesthetised and transcardially perfused with 4% PFA in PBS (see the section 4.2.2.) at four time points throughout 24 h: ZT0 (n=5), ZT6 (n=5), ZT12 (n=6), and ZT18 (n=6). Brains were extracted from the skull, post-fixed and cut on the vibroslicer in 40 μm thick sections. Slices containing the SC were immunostained against OXB and parvalbumin (PVA) with primary: goat-anti OXB and rabbit-anti PVA (1:1000; Abcam), and secondary antisera: donkey anti-goat Alexa Fluor 637 and donkey anti-rabbit Cy3 (1:300; Jackson ImmunoResearch). We used more efficient OXB antisera (in comparison to these against OXA) to visualise orexinergic cells and fibres at general, as OXA and OXB derive from a common precursor preproorexin and are co-stored [4]. Mounted sections were imaged under the confocal microscope under a 20× objective in 3 μm Z-stack steps.

### 4.4. Retrograde tract tracing

#### 4.4.1. Surgery

Neuronal tract tracing was performed on three Sprague Dawley rats (250-350 g) according to a procedure described previously [18]. In brief, isoflurane-anesthetised rats were mounted in the Small Animal Stereotaxic System (SAS-4100; ASI Instruments, Warren, MI, USA) and the same anaesthetic agent was continuously infused through the mask. Retrograde tracers: FluoroRed and FluoroGreen (20 nl; Tombow Pencil Co., Tokyo, Japan) were injected bilaterally into the superficial layers of the SC with 1 μl Hamilton syringe connected to borosilicate glass micropipette made at a vertical puller (Narishige, Tokyo, Japan). FluoroRed injection was aimed at left and FluoroGreen at right hemisphere: anteroposterior (AP) = −7.0, lateromedial (LM) = ±1.0, dorsoventral (DV) = −3.5 mm from Bregma. After seven days, rats were subjected to the same surgical procedure, during which the colchicine (0.1 mg in 5 μl saline; Sigma) was injected into the lateral ventricle (AP = −0.7, LM = +1.8, DV = −4 mm from Bregma). Subsequently, 24 h after the procedure rats were re-anesthetised with sodium pentobarbital and perfused with 4% PFA in PBS.

#### 4.4.2. Immunostaining and imaging

Brains were extracted from the skull, post-fixed and cut on the vibroslicer into 40 μm thick coronal sections containing the lateral hypothalamus (every third slice was collected from approximately −3.5 to −1.7 mm from Bregma in the anteroposterior dimension) and into 100 μm thick coronal midbrain slices containing the injection site. Next, the LH sections were immunostained against OXB (see 2.2.2) and imaged on the epifluorescence microscope under 20× magnification (Axio Imager M2; Carl Zeiss, Oberkochen, Germany).

### 4.5. Electrophysiological recordings *ex vivo*

#### 4.5.1. Tissue preparation

In total, 20 Sprague Dawley rats were culled at four time points across the day and night, namely at ZT0, 6, 12, and 18 (n=5 each). Brains were immediately removed from the skull and immersed in the ice-cold preparation artificial cerebro-spinal fluid (ACSF), composed of (in mM): 5 NaHCO_3_, 3 KCl, 1.2 Na_2_HPO_4_, 2 CaCl_2_, 10 MgCl_2_·6H_2_O, 10 glucose, 125 sucrose and 0.01 g l^−1^ phenol red (Sigma); constantly oxygenated with carbogen (95% oxygen, 5% CO_2_). Subsequently, midbrain sections were cut into 250 μm thick acute slices in a chamber of a vibroslicer filled with the same solution. Next, slices containing the SC were transferred to warm (32°C) recording ACSF solution, comprising (in mM): 125 NaCl, 25 NaHCO_3_, 3 KCl, 1.2 Na_2_HPO_4_, 2 CaCl_2_, 2 MgCl_2_·6H_2_O, 5 glucose and 0.01 g l^−1^ phenol red. Slices were incubated for 1 h before being transferred to the recording chamber of the multi-electrode array (MEA) system (Multi Channel Systems GmbH, Reutlingen, Germany).

#### 4.5.2. Recording

Slices were placed above the electrodes, anchored and positioned such that recording electrodes were in contact with the superficial layers of the SC near the midline of the brain. In all experiments, 6 × 10 perforated MEAs (100 μm spacing; 60pMEA100/30iR-Ti; Multi Channel Systems GmbH) were used. During the recording, gentle suction was applied to ensure proper contact with the recording electrodes and signal stability. Prior to the start of the recording, always two hours after the cull, slices were allowed to settle for half an hour. Slices were continuously rinsed with fresh, carbogenated ACSF (2 ml min^−1^) heated to 32°C. All drugs were applied by bath perfusion. Signal was sampled at 20 kHz and acquired with the Multi Channel Experimenter software (Multi Channel Systems GmbH) [37].

#### 4.5.3. Drugs

All drugs were stored as 100× concentrated stocks and were diluted in a fresh ACSF prior each application to their working concentration: OXA (200 nM, Bachem, Bubendorf, Germany), OXA17-37 (1 μM, Bachem), TCS-OX2-29 (20 μM, Tocris, Bristol, UK) and bicuculline (20 μM, Tocris).

### 4.6. Hybridisation *in situ*

#### 4.6.1. Tissue preparation

Six adult Sprague Dawley rats were deeply anaesthetised with isoflurane and culled by decapitation (n=3 at ZT6, n=3 at ZT18). Brains were then quickly removed from the skull, flash frozen upon the dry ice and stored at −80°C. Then, they were cryo-sectioned at −20°C on a cryostat (Leica CM1950), thaw-mounted on Superfrost-Plus slides (Fisher Scientific, USA) and stored at −80°C overnight. At the next day, sections were fixed in cold (4°C) 4% PFA solution in PBS for 15 mins, rinsed in fresh PBS and dehydrated in increasing ethanol concentrations (50, 70, 100, 100%). Following, slides were let to air-dry and each slice was drawn around with a hydrophobic barrier pen.

#### 4.6.2. RNAscope assay and imaging

Immediately following the tissue preparation, slices were processed with the RNAscope multiplex *in situ* hybridisation protocol (Advanced Cell Diagnostics – ACD, USA). First, slices were pre-treated with protease IV for 20 mins, rinsed in PBS and incubated for 2 h at 40°C with probes targeting *Hcrtr2*, *Per2* and *Penk*. Following, sections were rinsed with wash buffer and transferred to a four-step signal amplification protocol, terminating with the fluorophore tagging (*Hcrtr2* with Alexa 488, *Per2* with Atto 550, and *Penk* with Atto 647). Last, sections were rinsed twice with the wash buffer, stained with DAPI and coverslipped with the fluorescent mounting medium (ProLong™ Gold antifade reagent, Invitrogen, USA). Two LGNs per animal were imaged under 20 × magnification with the epifluorescence microscope (Axio Imager.M2, Zeiss, Germany) and images were inspected in ZEN software (ZEN 2.3. blue edition, Zeiss).

### 4.7. Real-time quantitative reverse transcription polymerase chain reaction (RT-qPCR)

#### 4.7.1. Animals and tissue preparation

Two cohorts of Sprague Dawley rats (24 each) were used in this study. First one was kept under standard 12:12 light-dark cycle (LD), and another was moved to the constant darkness (DD) 48 h before cull. Rats were culled in four daily/circadian time points across 24 h: ZT/CT0, 6, 12, and 18 (n=6/time point/light conditions). Brains were quickly extracted from the skull and placed in the cold preparation ACSF. Subsequently, they followed the same slicing procedure into 250 μm thick sections as described in 4.5.1. The superficial layers of the bilateral SC were then dissected from these midbrain slices with a surgical scalpel and immediately flash frozen upon the dry ice. These fragments of tissue were stored at −80°C up to one week. All instruments used for the preparation were surface treated with RNaseZap (Sigma), to minimise ribonuclease activity.

#### 4.7.2 RNA isolation and real-time RT-qPCR

RNA was extracted from the dissected fragments of tissue with the use of ReliaPrep RNA Tissue Miniprep System (Promega, Madison, WI, USA). The obtained RNA was stored in RNase-free water at −80°C. Next, reverse transcription was performed with the High-Capacity RNA-to-cDNA Kit (Applied Biosystems, Foster City, CA, USA), for the same amount of RNA from each sample. Real-time RT-qPCR was carried out with the PowerUp SYBR Green Mastermix (ThermoFisher Scientific, Vilnius, Lithuania) at the StepOnePlus Real-Time PCR System (Applied Biosystems). For transcript amplification, the QuantiTect primer assay was used (Qiagen, Hilden, Germany). The studied genes included: *Hcrtr1, Hcrtr2* (for OX_1_ and OX_2_ receptor, respectively)*, Arntl* (for Bmal1), *Nr1d1* (for Rev-Erbα), *Per1* and *Clock*, whereas *Gapdh* served as a housekeeping gene.

### 4.8. PER2∷LUC bioluminescence monitoring of slice explant

Four 11 week old male heterozygous PER2∷LUC mice (RRID: IMSR_JAX:006852) were used for bioluminescence monitoring experiments. Coronal brain slices containing anterior and posterior sections of the SC were prepared at 300 μm thickness in ice-cold HBSS (Gibco/Thermo Fisher Scientific, Waltham, MA) on a vibratome (Leica VT1000S, Heidelberg, Germany). The two slices respectively corresponded to the 26^th^ and the 28^th^ sections in the OX_2_ receptor expression data of the Allen Brain Atlas (ISH data: Hcrtr2 - RP_080424_01_A11 - coronal). The SC explants were quickly cut on a stereomicroscope and transferred to culture membranes (Millipore Millicell-CM, Bedford, MA) and cultured in a phenol-red-free Dulbecco’s modified eagle medium (DMEM) (Sigma) containing 2% B-27 supplement (Gibco), 4.2 mM sodium bicarbonate (Gibco), 10 mM HEPES (Gibco), 1% penicillin-streptomycin (Gibco), and 300 μM beetle luciferin (Promega, Madison, WI) in 35 mm dish sealed with silicone vacuum grease (Dow Corning, Midland, MI).

Bioluminescence from the PER2∷LUC reporter was continuously recorded from the culture with the LumiCycle (Actimetrics, Wilmette, IL), and later analysed with custom-written Mathematica package (PMTAnalysis.m) [38]. The raw PER2∷LUC time series was detrended by using the Hodrick-Prescott filter [38]. Spectral analysis was performed by sliding fast Fourier transform (FFT) with a sliding window of three days; the resulting spectrograms were therefore truncated by last three days. All time series and spectrogram presented are averages of independent data from four animals.

### 4.9. Data and statistical analysis

#### 4.9.1. Quantification of OXB-ir fibres

Regions of interest (ROIs) were identified with marker staining for PVA, and the OXB-ir was analysed for the superficial and intermediate layers separately. Images were binary converted and measured in Fiji software (NIH, USA) with the custom-made macro. First, Bernsen’s adaptive thresholding method was used to define regions of the high local contrast. Next, immunoreactive pixels representing OXB-ir fibres were counted in the six ROIs outlined by a 40×40-pixel oval for each SC layer, and then divided by the area for the measurement to represent the area fraction. Then, results from two slices for each rat were averaged. All images were analysed with the same settings. Statistical analysis was performed in Prism 7 (GraphPad Software, San Diego, CA, USA) with ordinary one-way ANOVAs.

#### 4.9.2. Cell counting

The LH slices were sorted from the most caudal to rostral and 14 slices of the same stereotactic coordinates were taken for further analysis from each brain. Cell counting was performed manually in ZEN software (ZEN 2.3. black edition; Zeiss, Germany). OXB-ir cells and these co-localising the retrograde dye were counted for each hemisphere. As every third slice was collected during the cutting procedure, all data were extrapolated by multiplying the results of cell counting by three. Results of overall cell counting are presented as average data from independent animals. The lateralisation of retrograde dye-filled OXB-ir cells relative to the injection side was statistically analysed in Prism 7 (GraphPad Software) with the paired t-test.

#### 4.9.3. Electrophysiology

##### 4.9.3.1. Spike sorting

Data were automatically spike-sorted in KiloSort program [39] working in the MatLab R2018a environment (MathWorks, Natick, MA, USA) as described before [18]. First, raw data were exported with Multi Channel DataManager (Multichannel Systems GmbH) to HDF5 files. These were further processed with a custom-made script in Matlab, for data remapping and conversion to DAT format. DAT files were first automatically spike-sorted with KiloSort, with a GPU was used to boost spike-sorting speed (NVIDIA GeForce GTX 1050Ti GPU; CUDA 9.0 for Windows). In parallel, raw data were exported with Multi Channel DataManager to CED-64 files. Following, they were filtered from 0.3 to 7.5 kHz (Butterworth band pass filter, fourth order) and remapped with custom-made Spike2 script (Spike2 8.11 software, Cambridge Electronic Design Ltd., Cambridge, UK). Finally, results of an automatic spike-sorting were transferred into the prepared CED-64 files. Then, manual refinement of spike-sorted putative single units was conducted in Spike2 8.11 with the use of autocorrelation, principal component analysis and evaluation of spike shapes.

##### 4.9.3.2. Analysis of firing rate and responses to drugs

Single unit data were next binned into firing rate histograms and further analysed in NeuroExplorer 6 (Nex Technologies, Colorado Springs, CO, USA). For the spontaneous firing rate analysis, data were 1800 s binned and compared in Prism 7 (GraphPad Software) with Kruskal-Wallis test. For the examination of drug response, data were 30 s binned. A unit was classified as activated by the treatment if the peak firing rate after the drug application exceeded three standard deviations (SDs) from the baseline mean. Similarly, if the single unit activity (SUA) was lowered by three SDs, the unit was deemed inhibited. Response amplitudes were calculated by extracting the 600 s baseline mean from the maximal/minimal firing rate value during the response. These were statistically compared in Prism 7 (GraphPad Software) with Friedman tests (for paired comparisons) or Kruskal-Wallis tests (for unpaired comparisons). Post-hoc testing of paired data was performed with Dunn’s multiple comparison test. Outliers were identified with ROUT test (Q = 0.01%). Pie charts were also generated in Prism 7 (GraphPad Software) and compared with Chi^2^ test.

##### 4.9.3.3. Heatmaps, average and bubble density plots

Heatmaps, average and bubble density plots were generated with the use of custom-made scripts in MatLab (MathWorks, Natick, MA, USA). For temporal heatmaps, data were 30 s binned and smoothed with the Gaussian filter (width: 5 bins). Next, the firing rate of each single unit was normalised from 0 to 1 to represent the minimal and maximal value throughout the recording. These were then sorted according to the relative SUA during the first response to OXA. Spatial heatmaps representing the amplitude of the response to OXA were calculated for each recording location by extracting the 400 s baseline multi-unit activity (MUA) from the mean 200 s MUA during the peak response. Average plots were generated based on the normalised SUA and presented as mean ± SEM. Bubble density plots and spatial density heatmaps were based on the location of single units, which position (a stereotactic coordinate in relation to Bregma) was extracted and classified into the distinct SC layer using the stereotaxic atlas of the rat brain [40]. Bubble density plots were colour-coded for excitation, inhibition and lack of response, whereas distinct special density heatmaps were generated for each condition. For the purpose of bubble density plot preparation, the surface containing positions of all units recorded was divided into small square subregions. Next, the number of units within each square was counted; for each square a bubble was drawn, with the size based on the number of units counted. Last, each bubble was repositioned to the centre of mass of all neurons located within the corresponding square.

#### 4.9.4. Relative quantification of gene expression

Data were analysed using the ΔΔCT method using *Gapdh* as a reference gen. Next relative target gene expression (RQ) was calculated with ZT/CT0 mean value as 1. For the comparison of *Hcrtr2* and *Hcrtr1* gene expression, RQ was calculated from the mean ΔΔCT for the *Hcrtr2*, which level served as a reference for both transcripts. Data were statistically tested in Prism 7 (GraphPad Software) with the use of ordinary one-way ANOVAs (for daily/circadian comparisons) or paired t-test (for relative receptor levels).

## Acknowledgements

Authors would like to thank Marcelina Janik, PhD for her valuable help with RT-qPCR data analysis. We would also like to thank the Department of Physiology and Toxicology of Reproduction for the access to their laboratory equipment.

